# Contigs directed gene annotation (ConDiGA) for accurate protein sequence database construction in metaproteomics

**DOI:** 10.1101/2023.04.19.537311

**Authors:** Enhui Wu, Vijini Mallawaarachchi, Jinzhi Zhao, Yi Yang, Hebin Liu, Xiaoqing Wang, Chengpin Shen, Yu Lin, Liang Qiao

**Affiliations:** Department of Chemistry, Shanghai Stomatological Hospital, Fudan University, Shanghai 200000, China; School of Computing, College of Engineering and Computer Science, The Australian National University, Canberra ACT 2600, Australia; Flinders Accelerator for Microbiome Exploration, Flinders University, Bedford Park SA 5042, Australia; Shanghai Omicsolution Co., Ltd., Shanghai 200000, China

**Keywords:** Taxonomic annotation, Database construction pipeline, Metaproteomics, Metagenomics, Gut microbiota, Deep proteome coverage

## Abstract

Microbiota are closely associated to human health and disease. Metaproteomics can provide a direct means to identify microbial proteins in microbiota for compositional and functional characterization. However, in-depth and accurate metaproteomics is still limited due to the extreme complexity and high diversity of microbiota samples. One of the main challenges is constructing a protein sequence database that best fits the microbiota sample. Herein, we proposed an accurate taxonomic annotation pipeline from metagenomic data for deep metaproteomic coverage, namely contigs directed gene annotation (ConDiGA). We mixed 12 known bacterial species to derive a synthetic microbial community to benchmark metagenomic and metaproteomic pipelines. With the optimized taxonomic annotation strategy by ConDiGA, we built a protein sequence database from the metagenomic data for metaproteomic analysis and identified about 12,000 protein groups, which was very close to the result obtained with the reference proteome protein sequence database of the 12 species. We also demonstrated the practicability of the method in real fecal samples, achieved deep proteome coverage of human gut microbiome, and compared the function and taxonomy of gut microbiota at metagenomic level and metaproteomic level. Our study can tackle the current taxonomic annotation reliability problem in metagenomics-derived protein sequence database for metaproteomics. The unique dataset of metagenomic and the metaproteomic data of the 12 bacterial species is publicly available as a standard benchmarking sample for evaluating various analysis pipelines. The code of ConDiGA is open access at GitHub for the analysis of real microbiota samples.

## Background

The human body is composed of both human cells and many different microorganisms that inhabit in various body sites. Through host-microbiota interactions, the microbes are closely associated with various diseases, including luminal diseases, immune diseases, metabolic diseases, neurodegenerative diseases, etc.[1-4]. To understand the role of microorganisms in human health and disease, it is necessary to characterize the changes in composition as well as the functional dynamics of microbiota. With the development of next-generation sequencing (NGS) techniques, metagenomics has greatly facilitated the study of microbiota[5, 6]. While metagenomics provides the information of composition and functional potential of microbiome, the method cannot determine proteins actually expressed in the microbiome[7, 8]. Proteins, as the biomolecules performing various functions within organisms, should be identified and quantified directly for microbiota function study.

During the past years, mass spectrometry (MS)-based metaproteomics is emerging as a powerful approach to understanding the functions of microbial communities[9-13]. Compared to the traditional proteomics of a single organism or simple mixtures, the metaproteomics of microbiota faces challenges of high complexity and taxonomic diversity, where dozens or even hundreds of species can be present in a sample with wide dynamic range variations, making the proteomic analysis extremely difficult[7, 14]. In shotgun proteomics, peptide identification relies on matching tandem mass spectra with protein sequences in a database. A crucial step in metaproteomics is to choose a suitable protein sequence database. An incomplete database can result in the missing of some key proteins, while a database with too large search space can result in limited detection sensitivity and high false discovery rates[15-17].

To date, the database building strategies in metaproteomics include mainly i) refining public proteome sequence databases, e.g. NCBInr[18] or UniProtKB/Swiss-Prot[19], using mass spectrometric data[20-24], ii) filtering public proteome sequence databases using taxonomic information by 16S rRNA sequencing[17, 25, 26], and iii) constructing sample-specific proteome sequence databases from whole-genome metagenomics sequencing[14, 16, 17]. Among the different strategies, sample-specific database construction by metagenomics sequencing has been considered as the best choice and employed in many metaproteomic studies[16, 27-30], wherein the databases contain only protein sequences specific to samples and hence can offer the best-fit search space to explore the protein expression of a particular microbiota and to improve the overall proteome coverage. However, it remains a challenging task to accurately construct an appropriate protein sequence database from metagenomic data due to potential issues in genome assembly, taxonomic profiling, etc.[14, 30]. Specifically, mis-assemblies of metagenomic contigs may lead to low accuracy on species-level annotation, while the relatively short lengths of genes may lead to low recall on gene-level annotations[17, 31]. Different approaches for taxonomic annotation of genes result in significantly divergent results in the downstream metaproteomic analyses[16, 28]. There is, to date, no consensus on a robust and reliable approach for constructing protein sequence databases, which limits the widespread use of metagenomic data for metaproteomic studies.

In this study, we developed an accurate taxonomic annotation strategy for metagenomic data to construct protein sequence databases, namely contigs directed gene annotation (ConDiGA). We mixed 12 known bacterial species to derive a synthetic microbial community to benchmark metagenomic and metaproteomic pipelines (**Figure 1**). Metagenomic sequencing-derived protein sequence databases with different taxonomic annotation strategies were compared. The optimal protein sequence database construction pipeline of ConDiGA was further applied to a human stool sample, and the metaproteomic identification results demonstrated the effectiveness of our strategy in real metaproteome samples. Our study promotes the in-depth metaproteomic analysis of microbiome and provides a reference for protein sequence database construction based on metagenomic sequencing.

**Figure 1.**
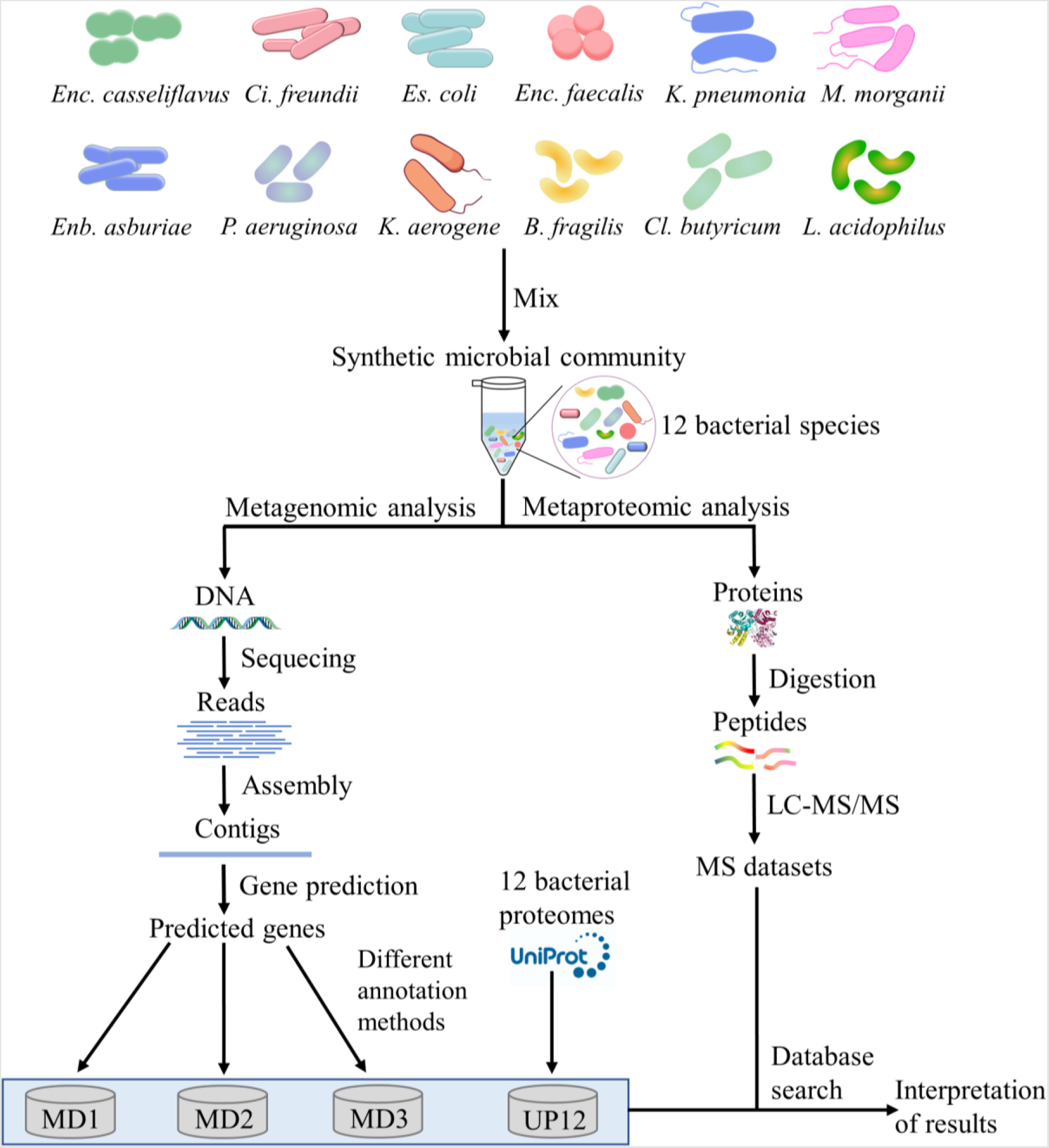
Benchmarking metagenomic and metaproteomic pipelines using a synthetic microbial community of 12 species. A synthetic microbial community with twelve species was constructed to conduct the comparative study of metaproteomics, and the optimization of protein sequence database construction from metagenomic sequencing data.

## Results

### Optimization of Protein Sequence Database Construction from Metagenomics Sequencing

A major difficulty in metaproteomics is the highly complex sample composition. How to optimize taxonomic annotation of genes in metagenomics is the key to generating an appropriate protein sequence database for metaproteomic analysis. We propose three different pipelines to construct three protein sequence databases (MD1, MD2, MD3, in **Figure 2A**). All the three pipelines first assembled raw reads into contigs using the metagenomics assembler MEGAHIT[32] (with the parameters k-min = 21 and k-max = 141) and then predicted genes from the assembled contigs using the gene identification tool MetaGeneMark[33]. Afterwards, three different taxonomic annotation strategies of genes were performed.

**Figure 2.**
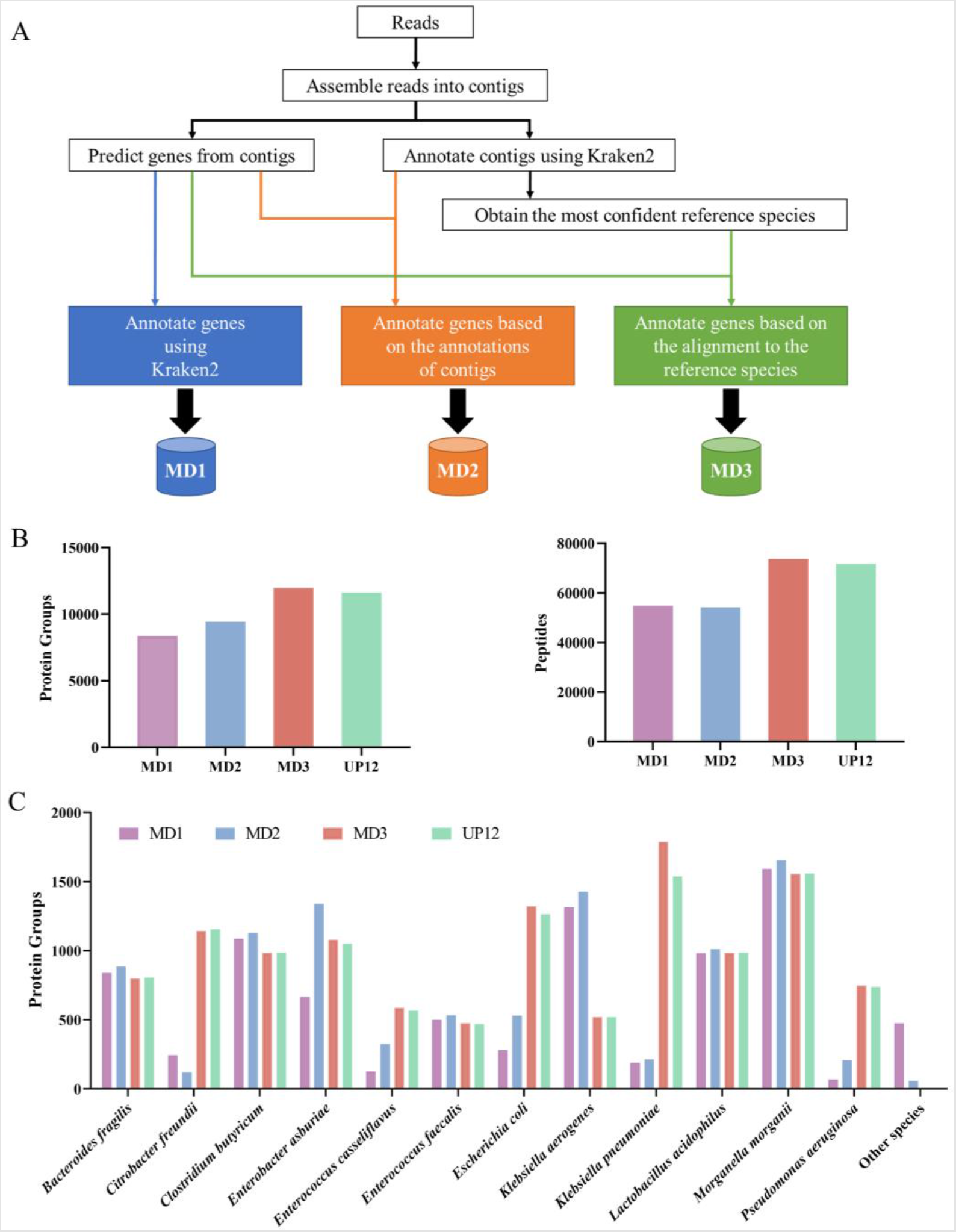
Optimization of protein sequence database construction from metagenomics. (**A**) The three pipelines MD1, MD2 and MD3 of constructing protein sequence databases from metagenomic data. (**B**) The number of protein groups and peptides identified using different databases. (**C**) The number of protein groups for each species identified using different databases. Two replications in LC-MS/MS analysis using TIMS-TOF PASEF were performed, and the combined identification results are shown. The sample was the synthetic microbial community of the 12 species.

In the first pipeline (MD1), the predicted genes were annotated using the popular taxonomic classification tool Kraken2[34] (with the parameter confidence = 0.1) against the whole bacterial genome database and the annotated genes were used to construct the MD1 database. As Kraken2 uses a k-mer based approach to individually annotate the predicted genes, it can introduce mistakes and result in a large number of annotated species which do not exist in the sample. To avoid this issue, the confidence parameter of Kraken2 was increased from 0 (default value) to 0.1. However, with the increase of the confidence parameter, many short genes cannot be annotated by Kraken2. Hence, the MD1 pipeline may result in a smaller number of annotated genes compared to the other databases.

In the second pipeline (MD2), the assembled contigs were annotated instead of predicted genes using Kraken2 (with confidence = 0.1). Then genes were annotated according to the species label of the contigs they belong to. As annotations on long contigs are more reliable than those on short genes, this pipeline can produce more annotated genes than the first pipeline. However, the species labels of contigs may be ambiguous or incorrect due to mis-assemblies, e.g., one mis-assembled contig might contain genes from multiple species (**Supplementary Figure 1**).

In the third pipeline (MD3), the assembled contigs were annotated using Kraken2 (with confidence = 0.1) and the most abundant species (with relative sequence abundance[35] > 0.5%) were selected. Then the predicted genes were annotated to one of the selected species based on the best alignment from Minimap2[36] to construct MD3. We name the third pipeline as contigs directed gene annotation (ConDiGA). In Latin, *condiga* means build, which well matches the process of building protein sequence databases from metagenomic data. Since the genes were annotated based on a refined set of species, MD3 not only has a higher recall on gene annotations but also is robust against misassemblies.

To benchmark the above three gene annotation pipelines, we mixed 12 known bacterial species to derive a synthetic microbial community (**Figure 1, Supplementary Table 1** and **Supplementary Table 2**). When applying this synthetic dataset, MD1 reports 48 species (0 false negatives and 36 false positives), MD2 reports 25 species (0 false negatives and 13 false positives), and only MD3 successfully recovers all the 12 ground-truth species (0 false negatives and 0 false positives). The detailed information in terms of the number of sequences and annotated species of the protein sequence databases built by the three pipelines are shown in **Supplementary Figure 2**.

We further compared the three protein sequence databases constructed by the MD1, MD2 and MD3 pipeline along with the reference protein sequence database from the 12 ground-truth species in Uniprot (named as UP12, **Supplementary Table 4**) on the analysis of metaproteomic data of the synthetic microbial community, where the data were acquired using the state-of-the-art MS technique of trapped ion mobility spectrometry (TIMS) synchronized with TOF for parallel accumulation serial fragmentation (PASEF) (TIMS-TOF PASEF, timsTOF Pro, Bruker). As shown in **Figure 2B**, the numbers of protein groups and peptides obtained using MD3 (11,975 protein groups) were the closest to and even a bit higher than those using UP12 (11,635 protein groups). By considering the number of identified protein groups and peptides, MD2 (9430 protein groups) was slightly better than MD1 (8353 protein groups), and MD3 was significantly better than both MD1 and MD2. As a further analysis, we compared the protein groups identified for each species using the 4 databases (**Figure 2C**). The numbers of protein groups assigned to each species using MD3 were basically consistent with those using UP12 except for *K. pneumonia*. The number of protein groups identified for *K. pneumonia* using MD3 was higher than that using UP12. UP12 database was constructed using the reference stains, which can be different from the strains in our synthetic microbial community. By metagenomic sequencing, it is possible to identify some proteoforms with sequence variations not included in UP12.

As for MD1 and MD2, there were significant differences in the number of identified protein groups for some species compared to the result by UP12 (**Figure 2C**). For example, the numbers of protein groups identified for *C. freundii, E. casseliflavus, E. coli, K. pneumonia* and *P. aeruginosa* using MD1 and MD2 were significantly smaller than those using UP12; while the numbers of protein groups identified for *K. aerogenes* using MD1 and MD2 were significantly larger than that using UP12. There were 475 and 57 protein groups identified for species not included in the synthetic microbial community using MD1 and MD2, respectively, which can be regarded as false identifications and accounted for 5.7% and 0.6% of the whole number of identified protein groups. The real false discovery rate (FDR) using MD1 was much larger than 1% although 1% FDR at both peptide and protein groups level was applied during database searching by PEAKS studio (Bioinformatics Solutions Inc.), demonstrating the significance of constructing a high-quality database with correct annotations. It should be noted that such error was introduced during database construction, especially the gene annotation, rather than the database searching, i.e., the protein sequences can be correct while the taxonomic annotations are wrong.

To check how the incorrect assignment was generated, identified proteins assigned to *K. aerogenes* and other species than the 12 species using MD1 and MD2 were searched against the protein sequences in the UP12 database using the basic local alignment search tool (BLAST, https://blast.ncbi.nlm.nih.gov/Blast.cgi). 199 protein groups identified for *K. aerogenes* using MD1 and 53 using MD2 were reassigned to *K. pneumonia*, respectively (**Supplementary Figure 3A** and **3B**). The protein groups assigned to other species using MD1 were mostly reassigned to *E. asburiae*; while most of the protein groups assigned to other species using MD2 were reassigned to *K. pneumonia* (**supplementary Figure 3C** and **3D**). The BLAST results can partially explain the significantly lower number of identified protein groups for *K. pneumonia* using MD1 and MD2, as well as the significantly lower number of identified protein groups for *E. asburiae* using MD1. It was then demonstrated that the errors were from the mistakes of gene annotation in MD1 and MD2, illustrating again the significance of gene annotation during the protein sequence database construction from metagenomic data.

As combined database strategies are often used in metaproteomics to increase the number of identifications, we merged the database MD3 with UP12 to get a non-redundant protein sequence database named UP12-MD3. A total of 56,250 protein sequences were included in the combined database. Identification results using the combined database were compared with those using the UP12 and MD3, respectively. As shown in **Supplementary Figure 4**, compared with MD3, the combined database UP12-MD3 did not lead to a much higher number of protein groups and peptide identification. This result highlighted the robustness and reliability of the MD3 database and hence MD3 is selected as the optimal database construction method. It is not recommended to merge extra databases into MD3, because the additional search space can lead to longer data processing periods and even lower sensitivity if the database is too large.

In addition to the optimization of the protein sequence database construction strategies, we also compared the state-of-the-art MS techniques of TIMS-TOF PASEF (timsTOF Pro, Bruker) and orbitrap (Orbitrap Fusion Lumos Tribrid mass spectrometer, Thermo Scientific) for metaproteomics using the above synthetic microbial community sample with a 2-hour data dependent acquisition (DDA) proteomics strategy. The timsTOF Pro was run with or without PASEF, and the latter was a typical Q-TOF strategy. To minimize the impact of the protein sequence database, the resulting MS datasets were then searched against the same reference database UP12. **Figure 3** illustrates the identification results with the different data acquisition strategies. Remarkably, the amount of protein groups and peptides identified with TIMS-TOF PASEF was significantly higher than those with Orbitrap and Q-TOF. The intersections among the protein groups identified with the three data acquisition techniques are illustrated as **Figure 3C**. 4589 protein groups were shared by all the three methods. 90% and 94% of protein groups identified by Orbitrap and Q-TOF were also identified by TIMS-TOF PASEF, and only 694 and 231 protein groups were unique to Orbitrap and Q-TOF, respectively. TIMS-TOF PASEF identified 52% and 125% more protein groups compared to Orbitrap and Q-TOF, respectively. The comparison of liquid chromatography coupled tandam mass spectrometry (LC-MS/MS) features and identified peptide-spectrum matches (PSM) showed similar results as the protein groups and peptides (**Supplementary Figure 5**), demonstrating an improved sensitivity and data capacity by TIMS-TOF PASEF under the same data acquisition time. The number of protein groups identified for each species was also counted (**Figure 3D**). TIMS-TOF PASEF obtained the highest identification number of protein groups for each species. The TIMS-TOF PASEF benefits the metaproteomic analysis on-one-hand by the enhanced data capacity and on-the-other-hand by the improved sensitivity. With TIMS, ions can be enriched according to their ion mobility and then released for MS analysis, resulting in an enhanced signal intensity. **Supplementary Figure 6** shows the MS1 signals of randomly selected precursors and their corresponding MS2 fragmentation spectra. With PASEF, the signal intensity as well as the number of matched fragments were clearly enhanced compared to those without PASEF.

**Figure 3.**
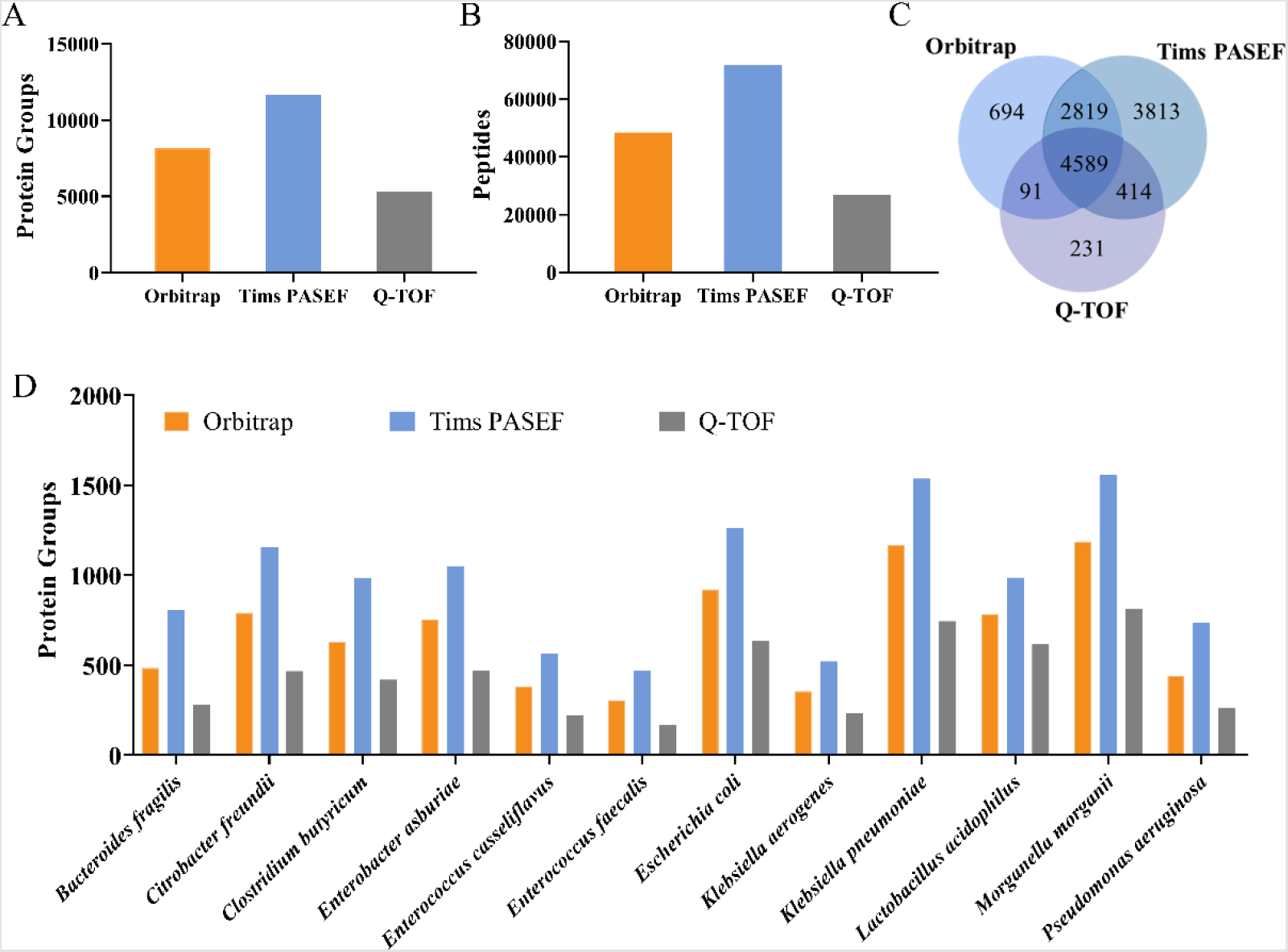
Metaproteomic analysis of the synthetic microbial community using Orbitrap, TIMS-TOF PASEF, and TIMS-TOF without PASEF (Q-TOF). (**A-B**) The number of protein groups and peptides identified by Orbitrap, TIMS-TOF PASEF and Q-TOF. (**C**) Venn diagram illustrating the overlap of protein groups identified by Orbitrap, TIMS-TOF PASEF and Q-TOF. (**D**) The number of protein groups for each species identified by Orbitrap, TIMS-TOF PASEF and Q-TOF. Three replications were performed for Orbitrap, and two replications were performed for TIMS-TOF PASEF and Q-TOF. The combined identification results are shown.

### Metaproteomic analysis of stool sample

To test the performance of the ConDiGA pipeline in real microbial communities, a stool sample from a healthy volunteer was collected and processed for metagenomic and metaproteomic analyses. We annotated the assembled contigs using Kraken2 (with confidence = 0.1) and then selected the species with a relative sequence abundance ≥ 0.01% and genome coverage ≥ 0.1%. A total of 77 species were selected for gene annotation under the criteria. Four protein sequence databases were constructed from the metagenomic data by considering the top 25 (TD25), top 50 (TD50), top 75 (TD75) most abundant, as well as all the 77 (TD77) species. The number of protein sequences of the databases are shown in **Supplementary Table 5. Figure 4A** show the protein identification results by TIMS-TOF PASEF with different databases. The numbers of identified protein groups and peptides using TD50 were significantly larger than those using TD25, but slightly smaller than those using TD75 and TD77. The highest number of peptide identification was reached using the TD75 database. When increasing further the number of species to 77, the search space was enlarged which led to a slightly lower number of identified peptides, while the number of identified protein groups were approximately the same. The numbers of protein groups for each species obtained using the four databases are shown in **Supplementary Figure 7**. *Faecalibacterium prausnitzii, Eubacterium eligens*, and *Phocaeicola vulgatus* were the top three abundant species identified using all the four databases, with more than 500 protein groups detected for each species. It was found that the TD25 can lead to a significantly higher number of proteins identified for species like *Phocaeicola vulgatus, Adlercreutzia equolifaciens*, and *Acutalibacter muris* compared to TD75 and TD77, which may be attributed to the wrong taxonomic annotation in TD25 due to the low number of species considered. Similar results were also observed for TD50.

**Figure 4.**
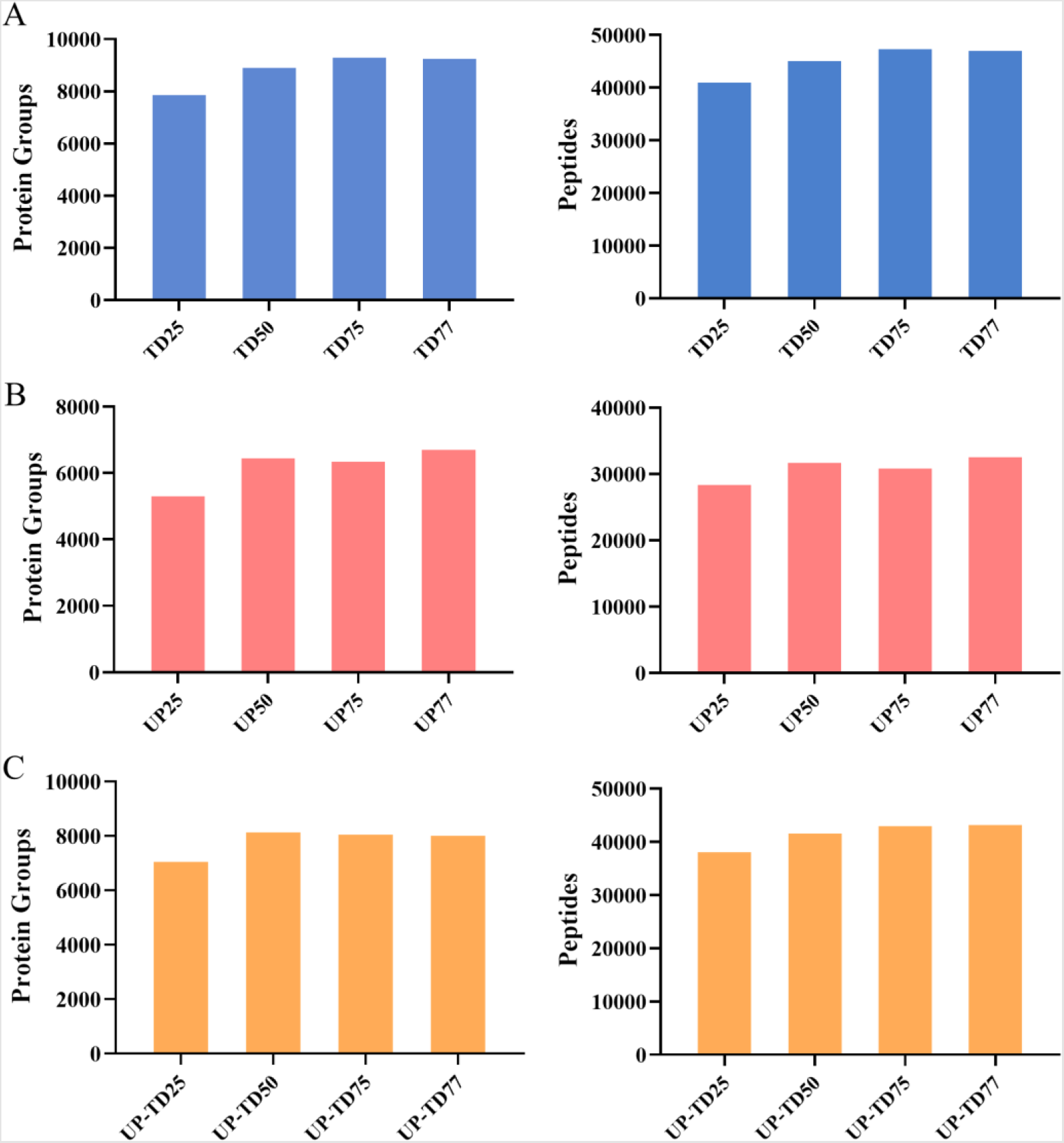
Metaproteomic analysis of a stool sample using different databases. The number of protein groups and peptides identified using different protein sequence databases constructed from (**A**) metagenomic data by ConDiGA pipeline, (**B**) Uniprot proteome databases of the corresponding species, and (**C**) the combined databases of Uniprot and metagenomics-based databases. Three replications in LC-MS/MS analysis using TIMS-TOF PASEF were performed, and the combined identification results are shown.

As a comparative analysis, databases were also constructed from the public proteome database (Uniprot) of the corresponding species (UP25, UP50, UP75, and UP77), as well as by combining the databases of Uniprot and metagenomics-derived databases (UP-TD25, UP-TD50, UP-TD75, and UP-TD77). **Figure 4B, Figure 4C** show the number of protein groups and peptides identified using these databases. The Uniprot-based databases performed much worse than the metagenomics-based databases. The combined databases also performed worse compared to the metagenomics-based databases. Gene detection rates of each species using TD77 and UP77 databases were comparatively shown in **Supplementary Figure 8**. Using TD77, the highest gene detection rate reached nearly 30%, and the gene detection rates of 21 species were > 10%. In contrast, the highest gene detection rate using UP77 was less than 20%, and only 4 species had a gene detection rate > 10%. It should be noted that the gene detection rates were calculated only considering the genes included in the protein sequence databases.

Considering all the results together, it is clear that the sample-specific database built by metagenomic sequencing performs was better than the taxonomically filtered Uniprot proteome database. The taxonomically filtered Uniprot database on-one-hand does not really match the sample by considering the strain-level variations and on-the-other-hand can contain many proteins not at a detectable level leading to an enlarged search space and hence restricted identification sensitivity. In the combined databases, although all the protein sequences in the metagenomics-based database were included, there was still the problem of the enlarged search space. Therefore, it is recommended to build sample-specific database by metagenomics. It is demonstrated that the ConDiGA pipeline for species annotation are applicable to real gut microbiota samples.

### Functional and taxonomic annotation at metagenomic and metaproteomic levels

To further analyze the stool sample, we conducted a comparative investigation of functional and taxonomic features at metagenomic (MG) and metaproteomic (MP) level. Metagenomic result with the ConDiGA pipeline and metaproteomic result using the TD77 database were annotated using KEGG Orthology (KO) terms by the GhostKOALA online service. **Figure 5A** shows the comparison of the KO functional annotation between MG and MP in terms of “metabolism”, “genetic information processing” and “environmental information processing”. Notably, the KO annotation results by the two meta-omics methodologies were mostly consistent. Carbohydrate metabolism, amino acid metabolism, and energy metabolism were the most abundant metabolism categories according to metagenomics and metaproteomics. Compared to metagenomics, metaproteomics enables deeper characterization of functional categories, and the term of “signaling molecules and interaction” was only revealed by metaproteomics. This result highlighted the divergence between metagenomics and metaproteomics in revealing functional potential of microbiota.

**Figure 5.**
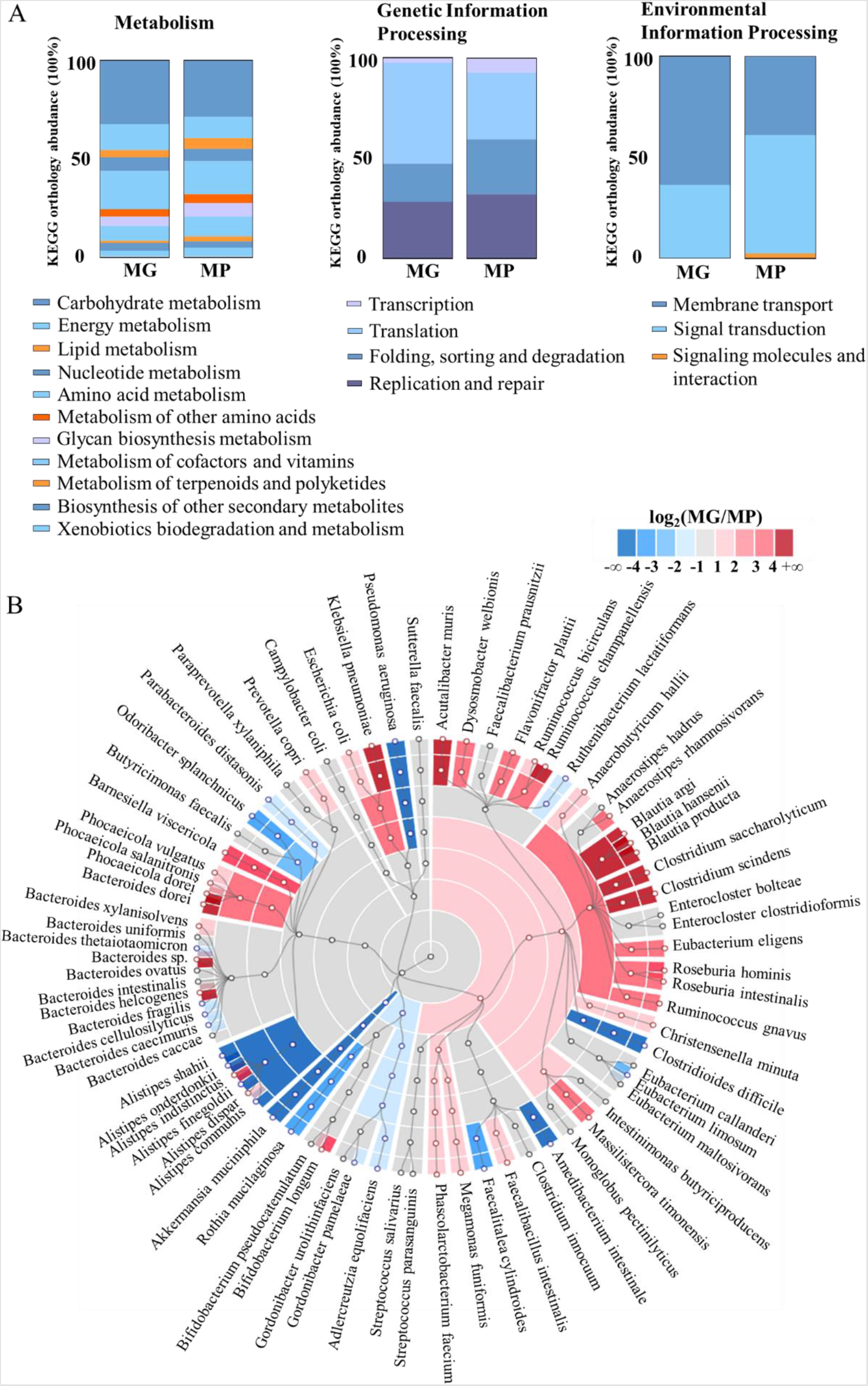
Comparative analysis of function and taxonomy at the metagenomic and metaproteomic levels. (**A**) KEGG functional annotation by metagenomics (MG) and metaproteomics (MP). The labels for the function terms are displaced in an order same as the stacked bar charts. (**B**) Taxa abundances by MG and MP. Colors indicate log_2_ ratio of relative abundance measured by MP/MG.

The relative taxonomic abundances at the metaproteomic level were computed by summing the quantitative information of all the identified peptides of each species, and the relative taxonomic abundances at the metagenomic level were calculated by counting the number of genes assigned to each species. The cladogram in **Figure 5B** depicts the relative abundance discrepancies between MP and MG by calculating the log MP/MG abundance ratio. The results showed that the abundance differences existed at different taxa levels. Five main phyla of bacteria, *Verrucomicrobia, Proteobacteria, Firmicutes, Bacteroidetes* and *Actinobacteria* were found in the stool sample, consisting with the previous studies[37]. Among them, *Firmicutes* and *Bacteroidetes* were more abundant by MG. At the species level, a total of 27 species showed higher abundance by MG, and 49 species were more abundant by MP. *Bacteroides dorei, Blautia argi, Ruminococcus champanellensis* and *Blautia producta* belonging to the phyla of *Bacteroidetes* and *Firmicutes* exhibited a significantly low MP/MG ratio. Interestingly, *Pseudomonas aeruginosa* (*Proteobacteria*), *Clostridioides difficile* (*Firmicutes*), and *Alistipes shahii* (*Bacteroidetes*) showed a significantly high MP/MG ratio. The results indicated that within the same phylum, the relative abundance of different species by metagenomics and metaproteomics can display heterogeneity. The relative abundance by MG is closely related to the cell copy of a species, while the relative abundance by MP shows the total protein amount of a species. Since different bacterial species can generate significantly different amounts of protein by one cell, it is reasonable that the relative abundances by MG and MP are different, and hence it is expected that MG and MP can show different sensitivity in the analysis of various species within a microbiota. Since proteins are the molecules performing various functions within organisms, it can be more accurate to predict functional potential of microbiota using the MP-based quantification information. The relative abundance discrepancies between MP and MG have also been reported by Tanca *et al*[37].

## Discussion

One of the most important issues of metaproteomics for complex microbial communities is the inadequate protein/peptide identification. To achieve a deep metaproteomic coverage of microbiota, robust MS measurements and efficient metaproteomic data interpretation pipelines are needed. The Bruker timsTOF pro can greatly increase sensitivity and reduce spectral complexity by synchronizing the trapped ion mobility spectrometry and time-of-flight mass spectrometry through the method TIMS-TOF PASEF. TIMS-TOF PASEF has been demonstrated as a powerful technique for deep proteomics on regular single organism or simple mixture samples, but its application in metaproteomics has not yet been reported. In this study, we explored the performance of TIMS-TOF PASEF in metaproteomics of microbiota. Utilizing TIMS-TOF PASEF, we achieved deep proteome coverage of about 12,000 proteins from the synthetic microbial community sample of 12 species by a 2-hour LC-MS/MS analysis. Compared to identifications from the same sample with PASEF disabled (Q-TOF) and Orbitrap, TIMS-TOF PASEF exemplified greatly expanded metaproteome coverage of microbiota.

It is generally recommended to use metagenomic data from the same sample to construct the protein sequence database for metaproteomic data analysis. Despite the fact that different metagenomics-based database construction strategies have been developed, an optimization of gene taxonomic annotation methods has not been reported, which, however, is extremely important for accurate metaproteomic results. In this study, we proposed three taxonomic annotation pipelines to construct metagenomics-based protein sequence databases and assessed their performance in metaproteomics using a lab-assembled microbial mixture as the benchmark sample. To date, one widely used approach in constructing metagenomics-based protein sequence databases is based on annotation on predicted individual genes [16, 28, 30] (MD1 in this study). We found that the popular pipeline could result in a large number of annotated species that are not really present in the sample due to inaccurate annotation on short genes. As annotations on long contigs are expected to be more reliable than those on short genes, we further constructed the second database (MD2) where the taxonomy information of contigs was passed on to its genes. The risk of this pipeline is that assembled contigs may suffer from misassemblies, resulting in incorrect annotations at the gene level. Therefore, we proposed the third annotation pipeline, named ConDiGA, which first selected the most abundant species from the annotation results of contigs, and then performed the gene-level annotation against these most abundant species.

## Conclusions

In summary, we achieved deep metaproteomic coverage of microbiota by developing an optimized protein sequence database construction strategy from metagenomics. We demonstrated the importance of correct gene annotation in constructing protein sequence database for metaproteomics and showed that metaproteomics can provide additional valuable information on function and taxonomy of microbiota compared to metagenomics. Our optimized taxonomic annotation pipeline can tackle the current annotation reliability problem in metagenomics-derived protein sequence database and can promote the development of metaproteomics. The metagenomic and the metaproteomic data of the 12 species benchmark sample and the codes of ConDiGA are open access, which can be used for the evaluation of various data analysis pipelines as well as the analysis of real microbiota samples.

