## Supplementary Information for "Contigs directed gene annotation (ConDiGA) for accurate protein sequence database construction in metaproteomics"

Wu et al.

**Supplementary Table 1.** The sources and cultivation conditions of 12 bacterial species.

| Strain | Source | Cultivation condition |
|---|---|---|
| *Escherichia coli* ATCC 25922 | ATCC | TSB 37°C, shaking, aerobic |
| *Citrobacter freundii* CICC 10404 | CICC | TSB 37°C, shaking, aerobic |
| *Enterococcus casseliflavus* ATCC 700327 | ATCC | BHI 37°C, shaking, aerobic |
| *Enterococcus faecalis* ATCC 19433 | ATCC | BHI 37°C, shaking, aerobic |
| *Pseudomonas aeruginosa* ATCC 27853 | ATCC | TSB 37°C, shaking, aerobic |
| *Enterobacter asburiae* ATCC 35953 | ATCC | TSB 30°C, shaking, aerobic |
| *Klebsiella aerogenes* ATCC 13048 | ATCC | TSB 30°C, shaking, aerobic |
| *Klebsiella pneumonia* ATCC 4352 | ATCC | TSB 30°C, shaking, aerobic |
| *Morganella morganii* CICC 21517 | CICC | TSB 30°C, shaking, aerobic |
| *Bacteroides fragilis* ATCC 25285 | CICC | BHI 37°C, shaking, anaerobic |
| *Lactobacillus acidophilus* CICC 6074 | CICC | MRS 37°C, shaking, anaerobic |
| *Clostridium butyricum* ATCC 1939 | CICC | MRS 37°C, shaking, anaerobic |

ATCC: American Type Culture Collection; CICC: China Center of Industrial Culture Collection; TSB: tryptic soy broth; BHI: brain heart infusion; MRS: De Man, Rogosa and Sharpe agar.

**Supplementary Table 2.** The composition of the simulated communities with 12 bacterial species.

| Species | Cell number ($10^8$ CFU) |
|---|---|
| *Morganella morganii* | 20 |
| *Pseudomonas aeruginosa* | 4 |
| *Klebsiella pneumonia* | 20 |
| *Citrobacter freundii* | 4 |
| *Enterococcus faecalis* | 2 |
| *Klebsiella aerogenes* | 2 |
| *Bacteroides fragilis* | 2 |
| *Enterobacter asburiae* | 10 |
| *Enterococcus casseliflavus* | 4 |
| *Escherichia coli* | 1.32 |
| *Clostridium butyricum* | 0.067 |
| *Lactobacillus acidophilus* | 0.333 |

**Supplementary Table 3.** LC gradients.

| Orbitrap | | timsTOF pro | |
|---|---|---|---|
| Time (min) | B (%) | Time (min) | B (%) |
| 0 | 2 | 0 | 2 |
| 3 | 2 | 105 | 22 |
| 96 | 17 | 110 | 35 |
| 111 | 27 | 115 | 80 |
| 116 | 37 | 120 | 80 |
| 119 | 50 | | |
| 121 | 95 | | |
| 130 | 100 | | |

**Supplementary Table 4.** Details of the reference proteome of 12 bacterial species.

| Strain | Protein count | Proteome ID/ Organism ID |
|---|---|---|
| *Escherichia coli* ATCC 25922 | 5062 | UP000000558/83334 |
| *Citrobacter freundii* CICC 10404 | 5149 | UP000305402/ 546 |
| *Enterococcus casseliflavus* ATCC 700327 | 3112 | UP000012675/ 565655 |
| *Enterococcus faecalis* ATCC 19433 | 3240 | UP000001415/ 226185 |
| *Pseudomonas aeruginosa* ATCC 27853 | 5564 | UP000002438/ 208964 |
| *Enterobacter asburiae* ATCC 35953 | 5254 | UP000216915/ 61645 |
| *Klebsiella aerogenes* ATCC 13048 | 4909 | UP000008881/ 1028307 |
| *Klebsiella pneumonia* ATCC 4352 | 5126 | UP000000265/ 272620 |
| *Morganella morganii* CICC 21517 | 3510 | UP000011834/ 1124991 |
| *Bacteroides fragilis* ATCC 25285 | 4234 | UP000006731/ 272559 |
| *Lactobacillus acidophilus* CICC 6074 | 1859 | UP000006381/ 272621 |
| *Clostridium butyricum* ATCC 1939 | 4245 | UP000003081/ 632245 |

**Supplementary Table 5.** Characteristics of protein sequence databases derived from the metagenomic data of a stool sample by considering different number of most abundant species.

| Database | Number of genes annotated | Fraction of genes annotated * |
|---|---|---|
| TD25 | 108,484 | 0.2318 |
| TD50 | 164,103 | 0.3506 |
| TD75 | 175,324 | 0.3745 |
| TD77 | 175,804 | 0.3756 |

*: the ratio of annotated genes to all the predicted genes.

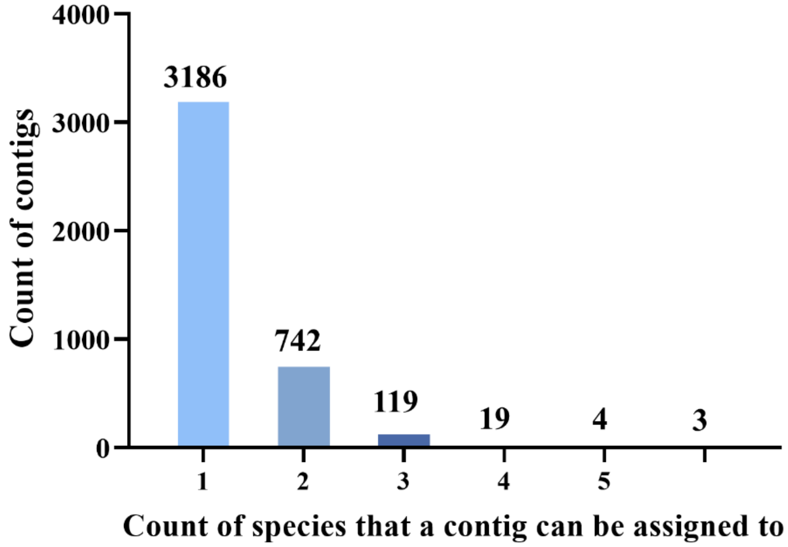


**Supplementary Figure 1.** Count of species in one contig assembled from the metagenomic sequencing results of the simulated microbial community of 12 species. There were 4073 contigs assembled. When at least 50% of a contig aligned to the genome of one species in the whole bacteria genome database, the species was identified from the contig.

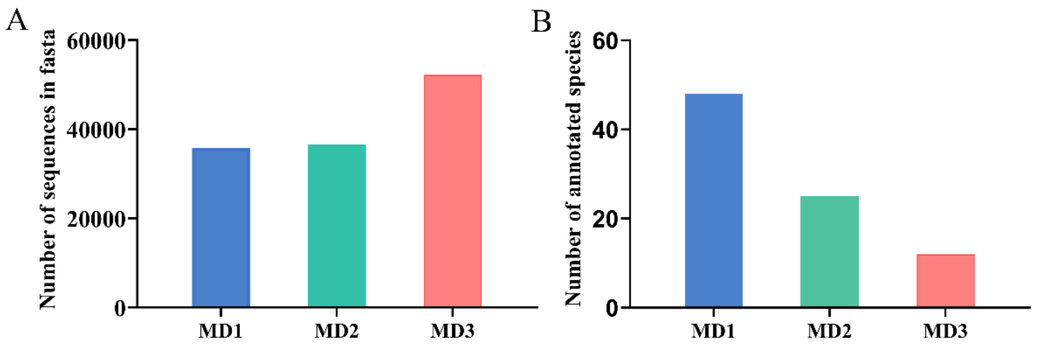


**Supplementary Figure 2. (A)** The number of annotated genes in the three metagenomics-derived databases for the simulated microbial community of 12 species. MD1:35,784, MD2: 36,582, MD3: 52,190. **(B)** The number of species annotated in the three metagenomics-derived databases for the simulated microbial community of 12 species. MD1: 48, MD2: 25 MD3: 12.

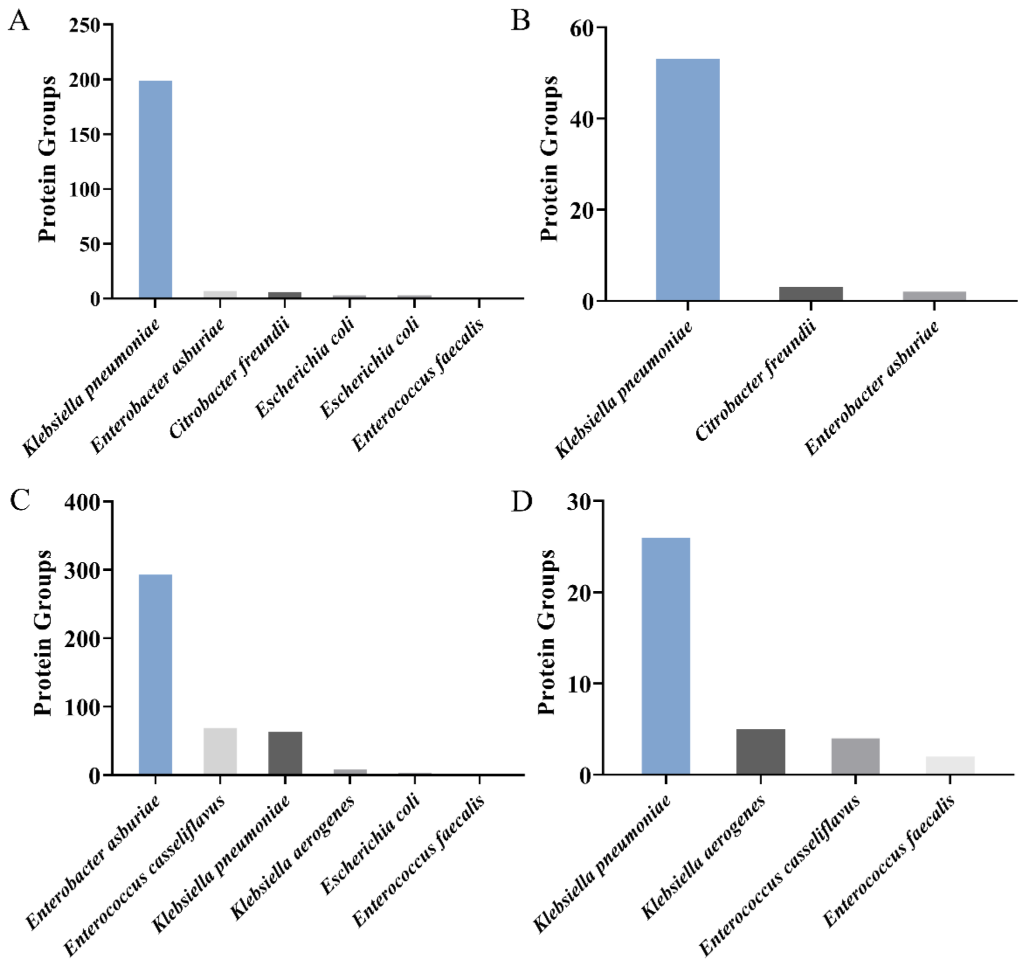


**Supplementary Figure 3.** BLAST results of the incorrectly assigned protein groups to *K. aerogenes* using **(A)** MD1 and **(B)** MD2. BLAST results of the protein groups incorrectly assigned to other species than the 12 species using **(C)** MD1 and **(D)** MD2. Sample: the simulated microbial community of 12 species; MS: TIMS-TOF PASEF. Two replications in LC-MS/MS analysis were performed, and the combined identification results are shown.

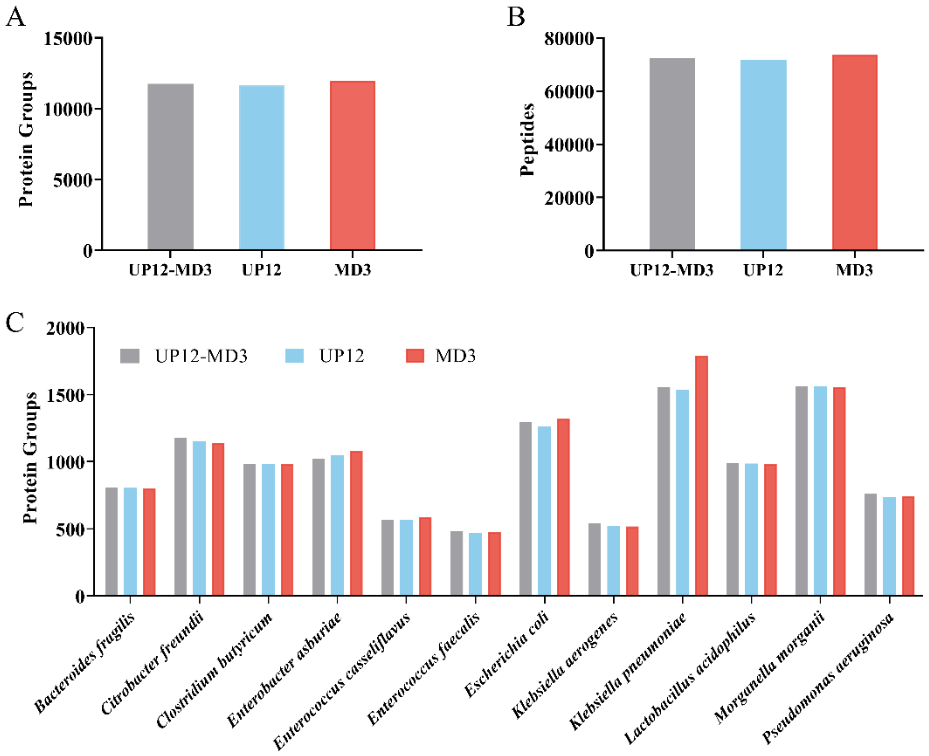


**Supplementary Figure 4.** Number of **(A)** protein groups and **(B)** peptides identified using UP12-MD3, UP12 and MD3. **(C)** Count of protein groups identified for each species using UP12-MD3, UP12 and MD3. Sample: the simulated microbial community of 12 species; MS: TIMS-TOF PASEF. Two replications in LC-MS/MS analysis were performed, and the combined identification results are shown.

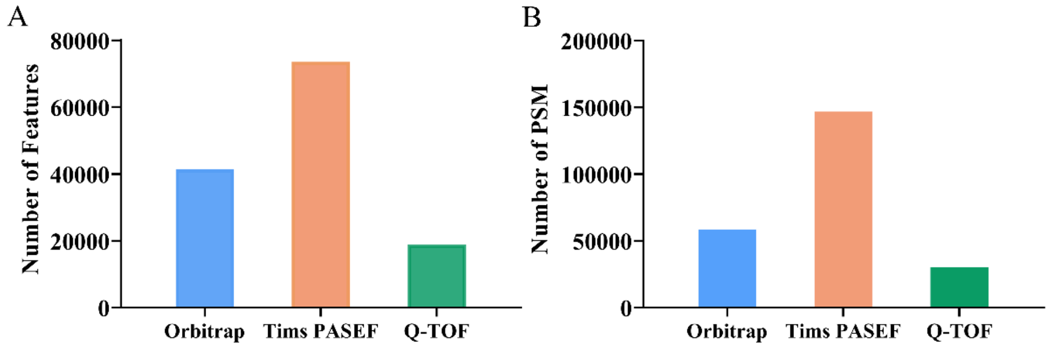


**Supplementary Figure 5.** The number of **(A)** Features and **(B)** PSM obtained by TIMS-TOF PASEF, Q-TOF, and Orbitrap from the simulated microbial community of 12 species. Three replications were performed for Orbitrap, and two replications were performed for TIMS-TOF PASEF and Q-TOF. The combined results are shown.

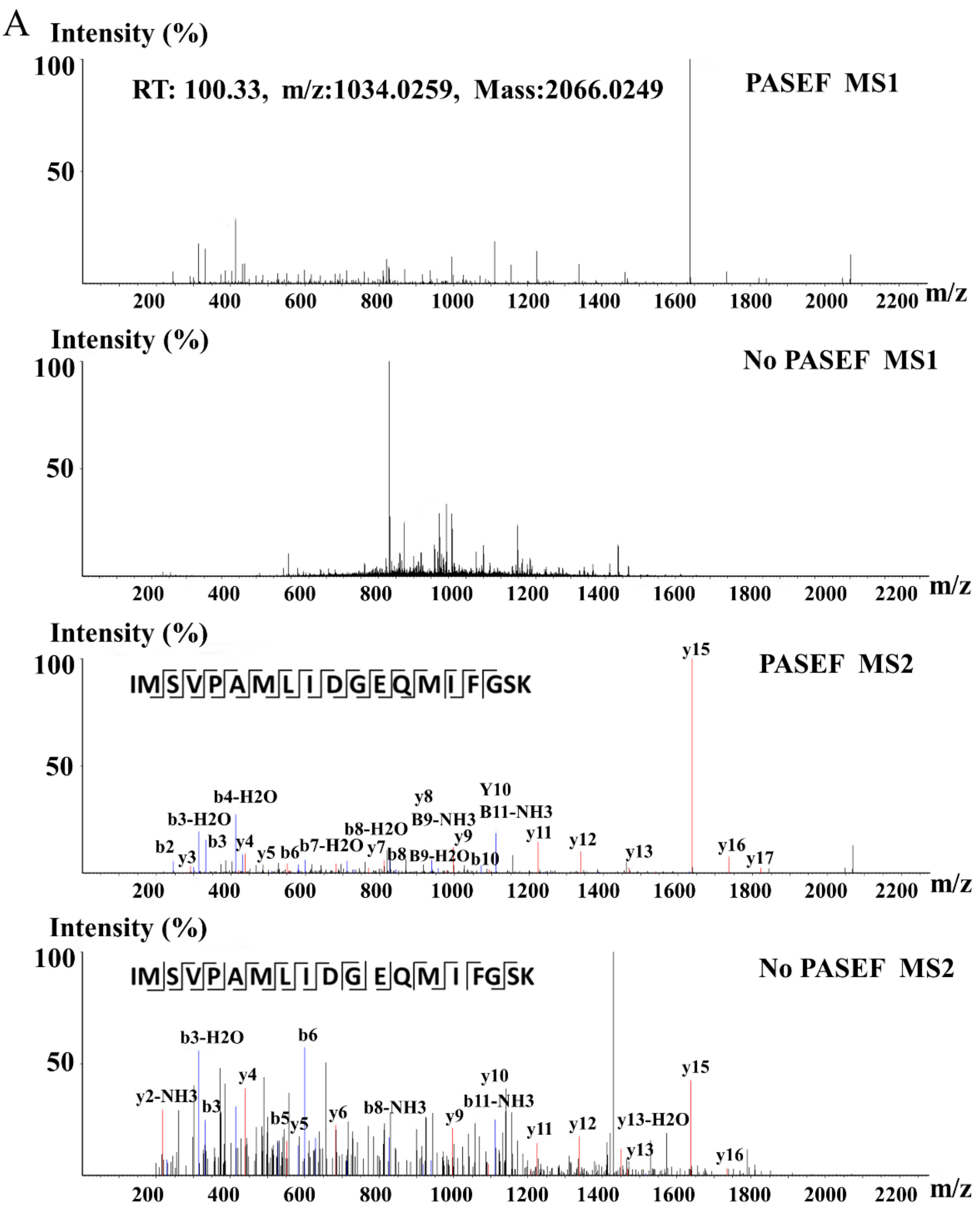

A
Intensity (%)
RT: 100.33, m/z:1034.0259, Mass:2066.0249
PASEF MS1
No PASEF MS1
PASEF MS2
IM|S|V|P|A|M|L|I|D|G|E|Q|M|I|F|GSK
No PASEF MS2
IM|S|V|P|A|M|L|I|D|G| E|Q|M| I |FG|SK
m/z

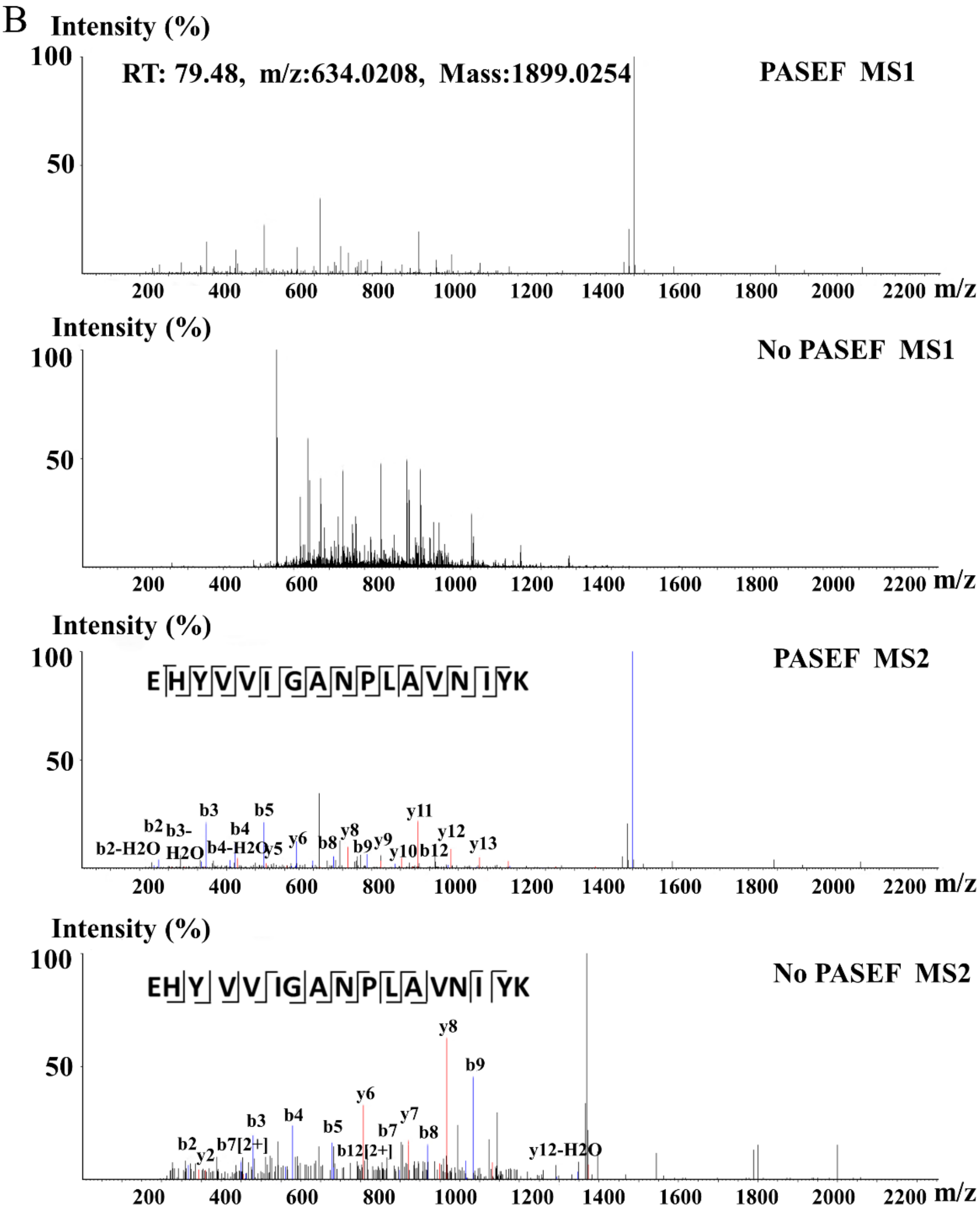


**Supplementary Figure 6.** Comparison of MS1 and MS2 signal of the same feature by timsTOF pro with and without PASEF when analyzing the simulated microbial community of 12 species.

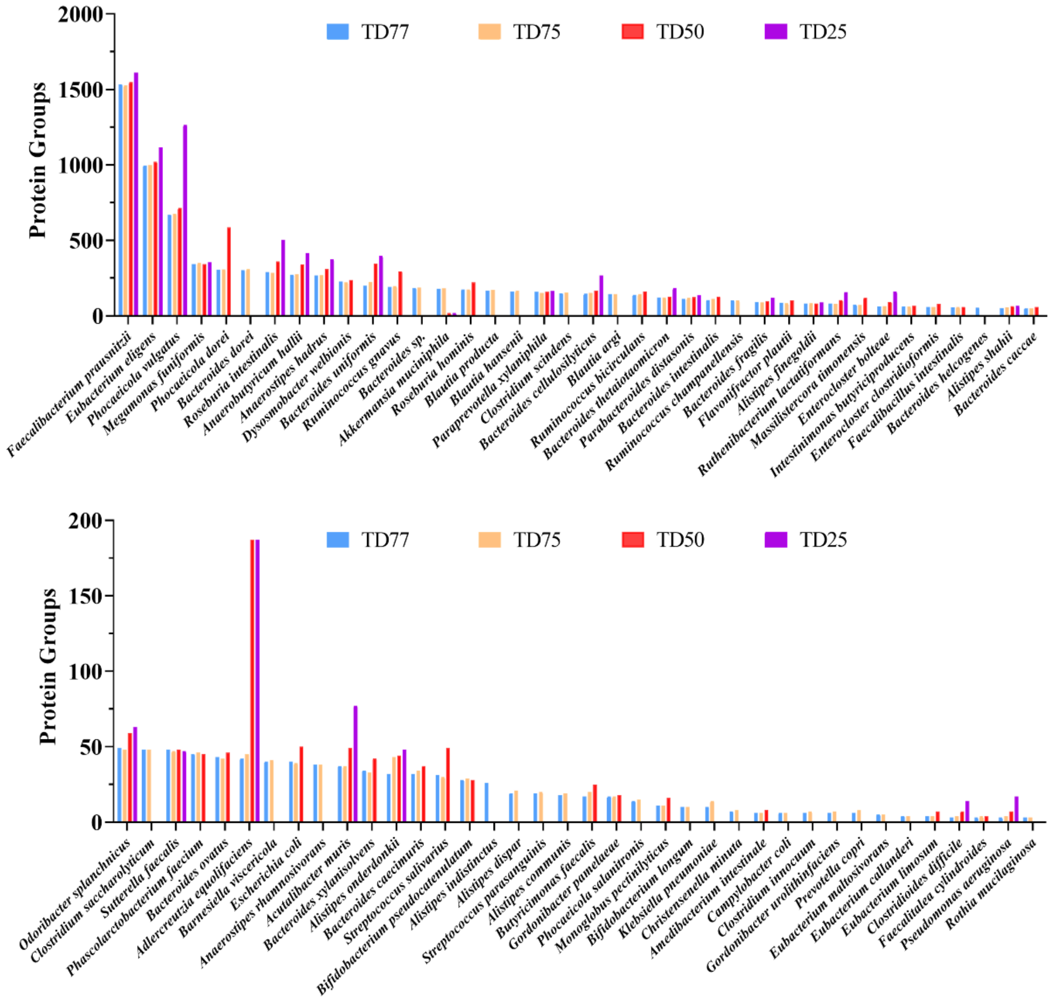


**Supplementary Figure 7.** Number of protein groups identified for each species from the stool sample using four metagenomics-derived protein sequence databases.

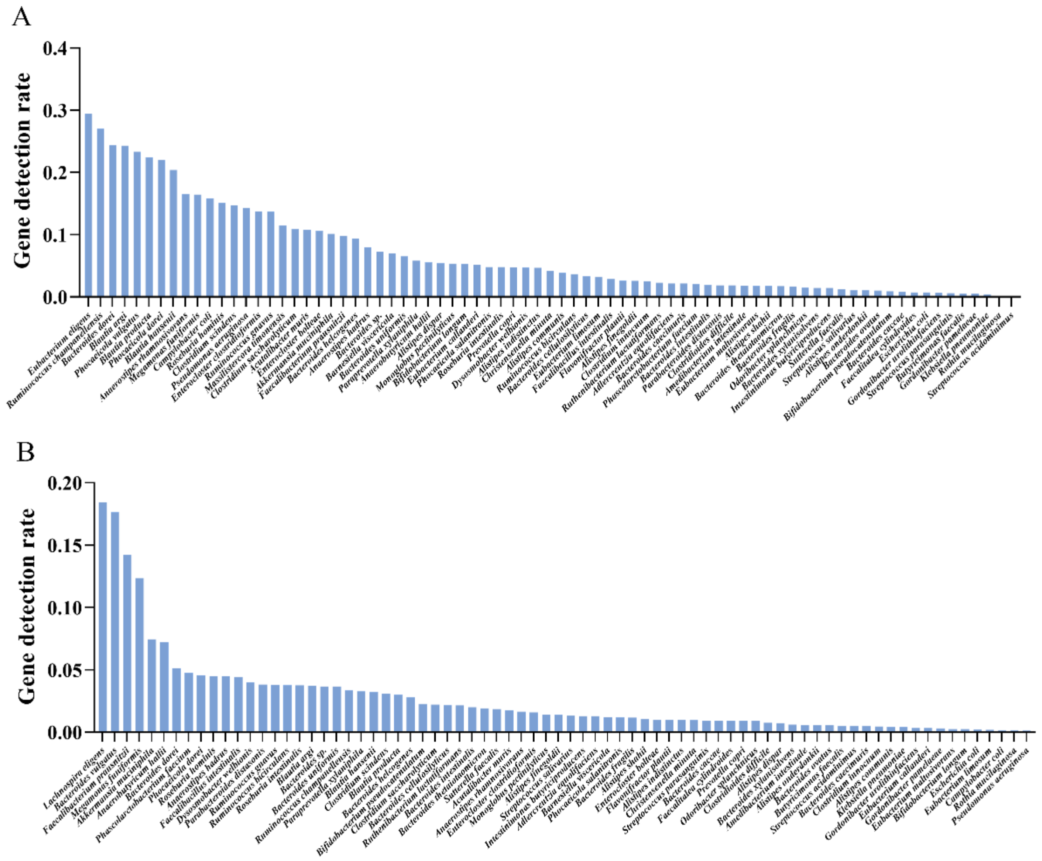


**Supplementary Figure 8.** Gene detection rates of each species from the stool sample using **(A)** TD77 and **(B)** UP77 databases.
